# Phage-encoded ribosomal protein S21 expression is linked to late stage phage replication

**DOI:** 10.1101/2021.10.11.463225

**Authors:** Lin-Xing Chen, Alexander L. Jaffe, Adair L. Borges, Petar I. Penev, Tara Colenbrander Nelson, Lesley A. Warren, Jillian F. Banfield

**Affiliations:** Department of Earth and Planetary Science, University of California, Berkeley, CA, USA; Innovative Genomics Institute, University of California, Berkeley, CA, USA; Department of Plant and Microbial Biology, University of California, Berkeley, CA, USA; Department of Civil and Mineral Engineering, University of Toronto, Toronto, Ontario, Canada; Department of Environmental Science, Policy, and Management, University of California, Berkeley, CA, USA; Chan Zuckerberg Biohub, San Francisco, CA, USA

**Keywords:** Phage, ribosomal protein, bS21, freshwater, Bacteroidetes virus

## Abstract

The ribosomal protein S21 (bS21) gene has been detected in diverse viruses with a large range of genome sizes, yet its *in situ* expression and potential significance have not been investigated. Here, we report five closely related clades of bacteriophages (phages) represented by 47 genomes (8 curated to completion and up to 331 kbp in length) that encode a bS21 gene. The bS21 gene is on the reverse strand within a conserved region that encodes the large terminase, major capsid protein, prohead protease, portal vertex proteins and some hypothetical proteins. These phages are predicted to infect Bacteroidetes species that inhabit a range of depths in freshwater lakes. Transcriptionally active bS21-encoding phages were sampled in the late-stage of replication, when core structural genes, bS21 and a neighboring gene of unknown function were highly expressed. Thus, our analyses suggest that bS21, which is involved in translation initiation, substitutes into the Bacteroidetes ribosomes and selects for phage transcripts during the late-stage replication when large-scale phage protein production is required for assembly of phage particles.

## Introduction

Only recently, ribosomal proteins have been recognized in the genomes of viruses ^1,2^, including those that infect bacteria (i.e., bacteriophages, or phages for short). Ribosomal protein S21 (bS21) is the most ubiquitous of the several ribosomal protein-encoding genes that have been reported in virus genomes ^1,2^ that range up to 642 kbp in length ^2^. In one study, viral bS21 was exclusively (over 90%) detected from aquatic samples ^1^. Functional assay experiments confirmed that the bS21 from Pelagibacter phage HTVC008M is incorporated into the 70S ribosomes in *Escherichia coli* ^1^, yet the *in situ* expression of bS21, and its potential significance to viral growth remain unclear. One hypothesis is that the viral bS21 protein will substitute for their host equivalent and may preferentially initiate translation of phage mRNA over bacterial mRNA ^2^. It has also been noted that phage bS21 homologs may contribute to specialized translation and/or help phages evade bacterial defenses ^3^.

The bS21 protein is small (8.5kD), highly basic, and specific for the bacterial ribosomes. It comprises two α-helices connected by a coiled region ^3^. It locates between the ‘head’ and the ‘body’ of the small ribosomal subunit (SSU) ^3^, in contact with the RNA helix formed between the mRNA and the 3’ terminus of the SSU ribosomal RNA (rRNA). This SSU region, also known as the anti-Shine-Dalgarno (ASD) sequence, is crucial for translational initiation by binding the mRNA Shine-Dalgarno (SD) sequence ^4^. Generally, when bS21 is missing, translation initiation is disturbed and the mRNA has lower association rates with the SSU ^5,6^.

In this study, we report 47 phage genomes that we assigned to five closely related clades of phages whose genomes consistently encode a copy of bS21. Notably, the bS21 gene colocates with genes for structural proteins that are responsible for virion assembly including the large terminase (TerL), portal vertex protein (PVP), prohead protease, and major capsid protein (MCP), but is encoded on the opposite strand. We manually curated all genomes and two outgroup phage genomes (thus 49 in total) to ensure accurate protein sequence prediction and to determine overall genome structure and genome sizes (when complete genomes were achieved). Nine genomes were completed, the largest of which is 331 kbp in length. CRISPR-Cas spacer targeting, taxonomic similarity of phage proteins to bacterial proteins, including bS21, all predicted that these phages infect freshwater Bacteroidetes species. We find that phage bS21 gene expression is significant during late-stage phage replication, likely specifically translating genes encoding core structural proteins that are essential to virion assembly and the lytic cycle.

## Results

### Discovery of closely related phage sequences with the conserved genetic context of bS21

Multiple phage-related sequences with a conserved genomic context were detected from several freshwater metagenome-assembled datasets (see Methods). Genes for bS21, terminase large subunit (TerL), portal vertex protein (PVP), prohead core scaffolding and protease protein (hereafter prohead protease for short), and major capsid protein (MCP) are encoded in the genomic region. BLASTp search of the TerL sequences against the ggKbase sequences (https://ggkbase.berkeley.edu) obtained a total of 47 unique scaffolds with the conserved genomic region (**Supplementary Table 1**). Two related phages were included as outgroups for comparative analyses.

### General features of manually curated genomes

All the 49 phage sequences were manually curated to fill scaffolding gaps and fix the assembly errors, and nine of them (including one outgroup phage) were curated to completion (circular and no gaps or local assembly errors) (**Supplementary Table 1**). A total of 14 related phage genomes from IMG/VR were also included for further analyses. The eight bS21-encoding complete genomes had genome lengths of 293-331 kbp, GC contents of 31.0-33.7% and encoded 350-413 protein-coding genes (coding density, 91.1-94.9%), with 5-25 (average 17) tRNA genes. No alternative coding signal (i.e., stop codon reassignment) was detected in any genome. In comparison, the outgroup complete genome has a size of 308 kbp (450 protein-coding genes, 6 tRNAs, 94.7% coding density) and GC content of 27.3%.

### Genomic context of bS21 in phages

Genomic context analyses for bS21 genes showed a highly conserved gene architecture across phage genomes in proximity to the region encoding bS21 (see **Figure 1a** for example). Specifically, we found that bS21 was consistently located in between two hypothetical protein families (positions 1 and −1 in **Figure 1b, Supplementary Table 2**), with core structural proteins - including the TerL, PVP, prohead protease, and MCP - generally located within 5 genes in both the upstream and downstream DNA. Other hypothetical proteins were also consistently found in this region, although their positions were more variable upstream (positions −4 through −10, **Figure 1b**). Importantly, the bS21 gene was consistently encoded in the reverse strand relative to the conserved hypothetical and structural protein genes (**Figure 1a, Supplementary Figure 1**).

**Figure 1.**
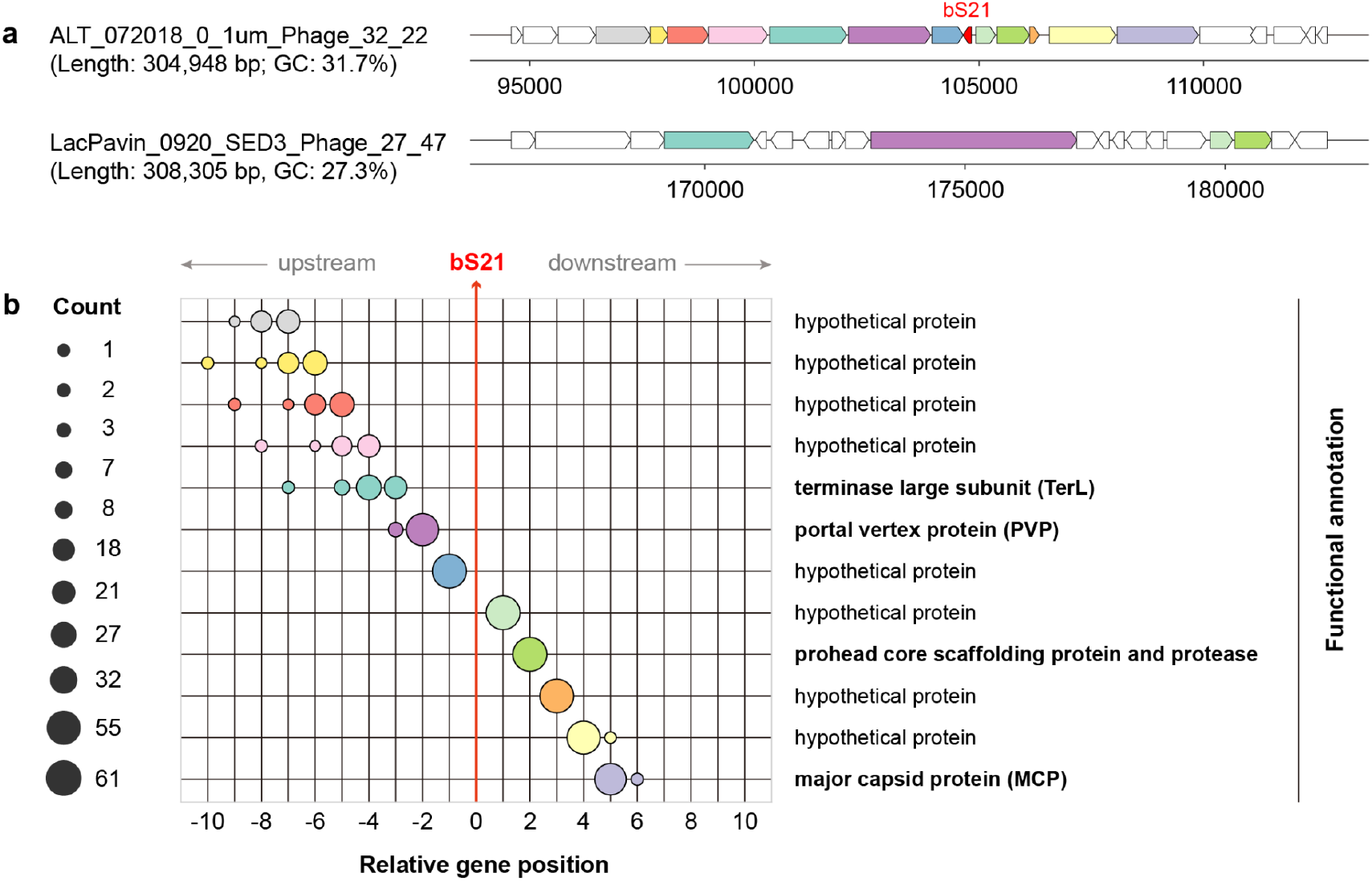
Genetic context of the genes encoding bS21 in the phage genomes. (a) Examples of genetic context of phage genomes with and without bS21. The annotation of protein-coding genes is the same as indicated in (b) by different colors. Those in white are genes not shown in subfigure (b). (b) Summary of genetic context of all phage genomes encoding bS21. The relative position of genes near the bS21 gene is shown, and the size of circles indicates the number of phages with a gene belonging to a given protein family (annotation shown on right) at that relative position. Only the 12 most frequent families are shown. The details of the genetic context are shown in **Supplementary Figure 1.**

### Phylogeny of bS21-encoding phages

Phylogenetic analyses based on TerL suggested the phages belonging to several groups, we thus assigned them to clades a-e (**Figure 2, Supplementary Table 1**). Most of the phages belong to clades c, d and e, and they have a broader environmental distribution than clades a and b. Interestingly, we found that some phages within a single clade were from distant sampling sites. Closer inspection indicated they also shared large genomic fragments with high similarity (82-98% for nucleotide sequences; **Supplementary Figure 2**). Comparative genome-wide analyses of the complete genomes from the same site but sampled at different time points showed sequence variations in some genes (**Supplementary Figure 3**).

**Figure 2.**
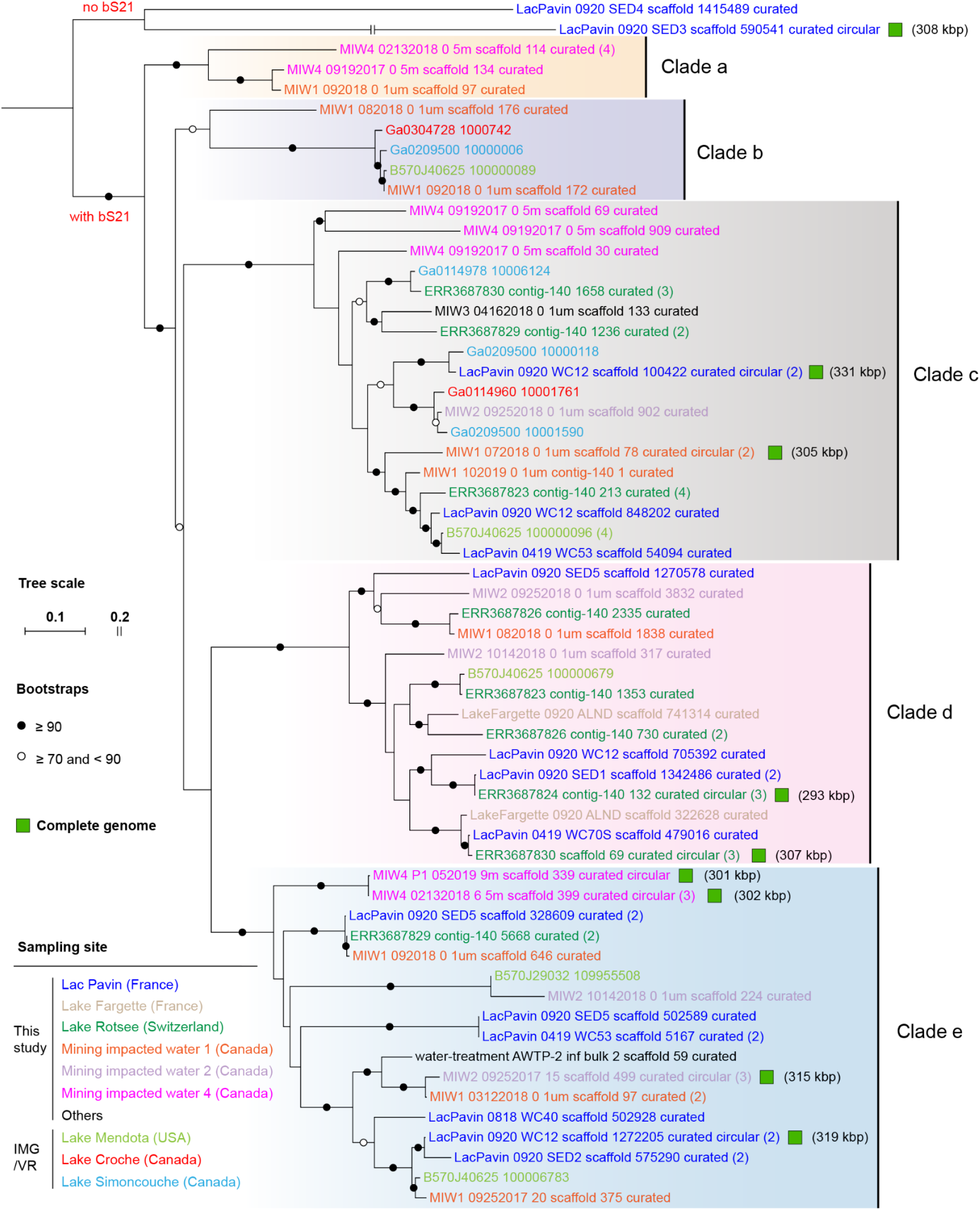
The phylogeny of bS21-phages based on the large terminal (TerL) protein sequences. Two closely related phages without bS21 encoded were included as outgroups (shown at the top of the tree). The genomes are assigned to five clades (a, b, c, d and e) based on the topology of the phylogenetic tree. The numbers in the brackets following the scaffold names indicate the total counts of the same scaffold detected from the corresponding sampling sites. The genomes that were manually curated to completion (circular and no gap) are indicated by squares, and the genome sizes are shown in brackets.

TerL phylogeny, constructed using sequences from this study and NCBI RefSeq sequences, indicated the most closely related classified phages belong to Caudovirales of either the Myoviridae or Ackermannviridae (**Supplementary Figure 4**). A phage baseplate assembly protein was encoded in most curated genomes. This is an important building block for members of Siphoviridae and Myoviridae ^7^, so we concluded that the bS21-encoding phages are myoviruses.

### Predicted bacterial hosts of bS21-encoding phages

To predict host-phage relationships we first used CRISPR-Cas spacers targeting. While none of the 16.5k unique spacers from the relevant metagenomes targeted any of the curated phage genomes from the same sampling sites, a single cross site target was detected. Specifically, MIW1_072018_0_1um_scaffold_78 was targeted by a spacer (24 nt and no mismatch) from a MIW2 *Flavobacterium* genome (affiliation: Bacteroidetes, Flavobacteria). We then predicted the bacterial hosts based on the bacterial taxonomic affiliations of the phage gene inventories as previously described ^2^ (**Supplementary Table 3**). The results indicated that all of the phages infect members of Bacteroidetes, which were detected in 43 out of 45 samples (**Figure 3, Supplementary Table 4**). The two metagenomic samples without Bacteroidetes identified were both collected via filtering through 0.2 μm and onto 0.1 μm pore size filters. Bacteroidetes were detected in both of the corresponding 0.2 μm fraction samples (**Figure 3**).

**Figure 3.**
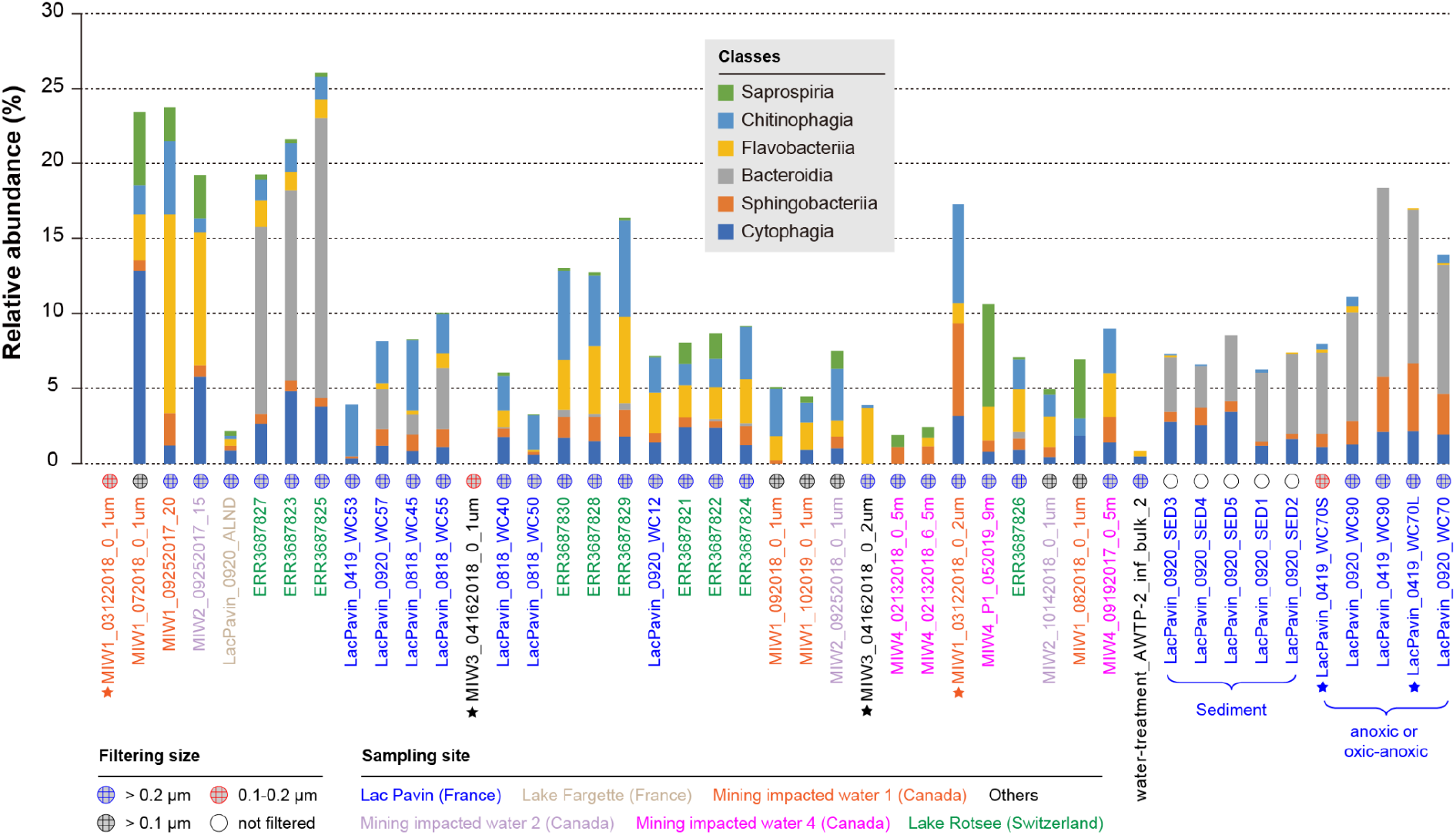
The relative abundance of the Bacteroidetes classes in all the analyzed samples in this study. The microbial communities were profiled based on ribosomal protein S3 (rpS3) assigned to the Bacteroidetes classes. The sampling sites were indicated by colored names, and the filter sizes used during sampling are shown by circles. The three pairs of filter samples are indicated by colored stars.

We profiled the co-detection of phage clades and Bacteroidetes classes to test for specific connection (**Supplementary Figure 5**). However, this was uninformative because most samples contained more than one class. However, phages from clades a and b are unlikely to infect class Bacteroidia members, as they did not co-occur in any sample.

### Comparison of bacterial and phage encoded bS21

Phylogenetic analyses revealed that bS21 protein sequences from phages (this study) and the bacterial bS21 sequences (from the corresponding samples and NCBI RefSeq) clustered separately (**Supplementary Figure 6**). The bacterial bS21 sequences that are most similar to phage bS21 were from Bacteroidetes, mostly from the Flavobacteriia class (**Supplementary Table 5**). We aligned and compared the Bacteroidetes and phage bS21 sequences and mapped the divergent and non-divergent residues to the model of the ribosome of *Flavobacterium johnsoniae* (**Figure 4a**). Multiple divergent positions are located at the beginning of the bS21 sequences and four residues (Arg21, Phe23, Asp25, and Thr28) were significantly divergent (**Figure 4b**).

**Figure 4.**
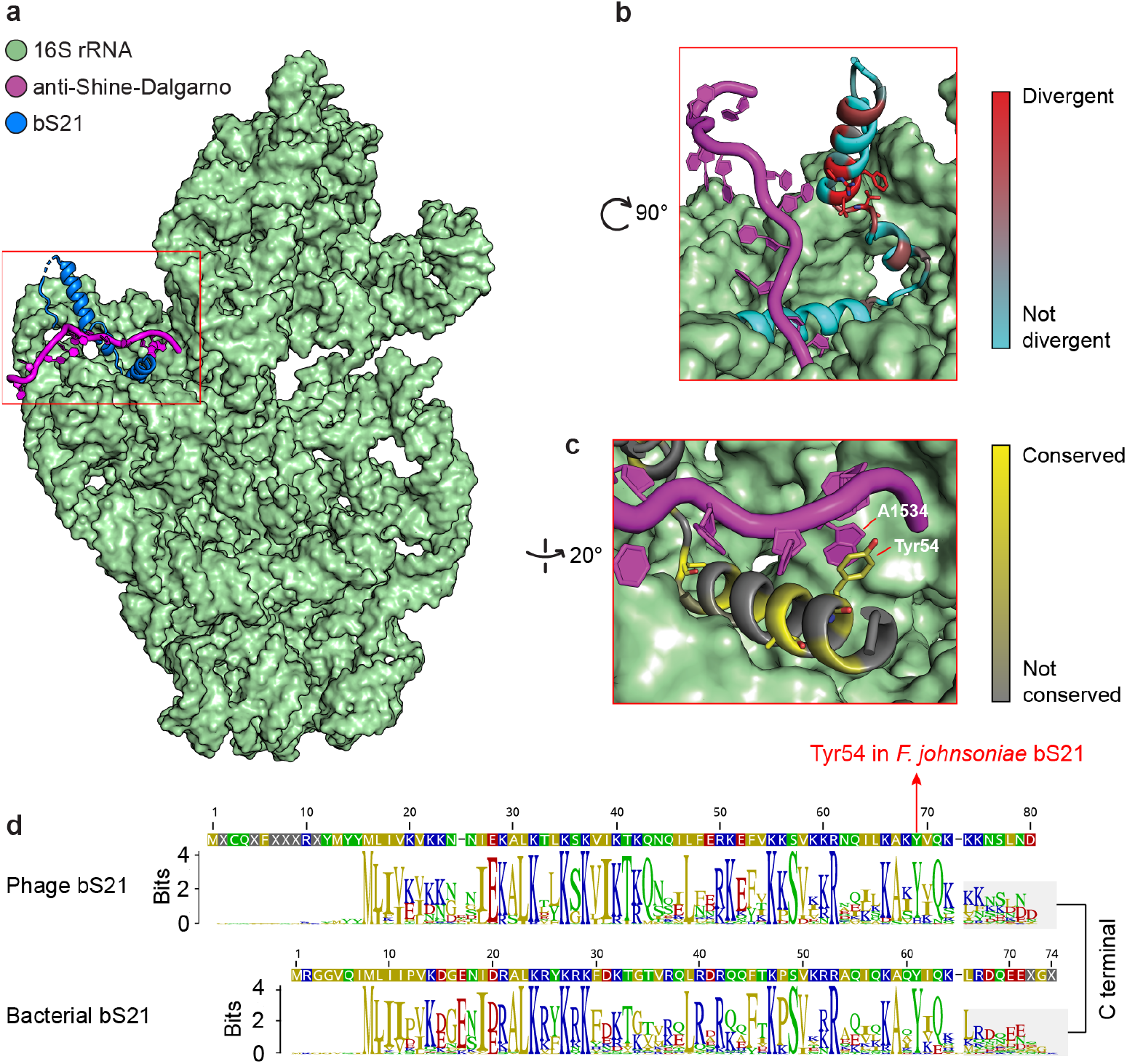
Conservation and differences between phage and bacterial bS21. (a) Location of bS21 (blue) within the 16S rRNA (green) and the ASD (magenta) of the *F. johnsoniae* ribosome (PDB ID: 7JIL) ^8^. bS21 is in the neck region of the 16S rRNA, interacting closely with the 3’ end of the 16S rRNA, where the ASD is located. The 16S rRNA is shown from the subunit interface direction. (b) Zebra2 divergency results from an alignment of phage and bacterial bS21 sequences mapped on *F. johnsoniae* bS21. Divergent positions between phage and bacterial bS21 are shown with red. (c) Zebra2 conservation results from the same alignment as in (b) mapped on *F. johnsoniae* bS21 with conserved residues shown in yellow. The stacking interaction between Tyr54 and Adenine 1534 is indicated. (d) The sequence logo and consensus sequences of phage and bacterial bS21 alignments, the corresponding position of Tyr54 in *F. johnsoniae* bS21 in the alignment is highlighted. The C terminal parts are highlighted with gray backgrounds.

Bacteroidetes usually lack the SD sequences. It was recently reported that the bS21 Tyr54 (numbering in *F. johnsoniae*) is an important residue for blocking the ASD in the 16S rRNA within the ribosome ^8^. Our analyses predict that all bacterial and phage bS21 have an amino acid with an aromatic ring (often Tyr54 but in a few cases His54, and in one case Phe54) at the position of Tyr54 in *F. johnsoniae* (**Figures 4c and d, Supplementary Figure 6**). This should ensure stacking interaction with Adenine 1534 (numbering in *F. johnsoniae* 16S) from the ASD.

In contrast, the C terminal regions of both the bacterial and phage bS21 sets were highly divergent (**Figure 4d**). However, the phage C terminal regions are generally conserved within the clades defined based on TerL phylogeny (**Figure 2, Supplementary Figure 7**).

### Metabolic potentials of bS21 encoding phages

Functional annotation of the predicted protein-coding genes revealed that in addition to bS21, these phages carry other genes related to protein production and stability (**Supplementary Table 6**). Examples include protein folding chaperones and Clp protease, suggesting the importance of controlling the proteostasis network of the cell. Interestingly, we also identified many genes involved in sugar-related chemistry and polysaccharide biosynthesis. Many of these genes were predicted to perform chemical transformations related to the biosynthesis of lipopolysaccharide, a major component of the Gram-negative bacterial outer membrane. We interpret this as a potential mechanism to remodel the cell surface and prevent superinfection by competitor phages, a strategy common to the phage lysogenic cycle. These phages lack detectable integration machinery (no gene for integrase or resolvase was detected), suggesting the possibility of a non-integrative long-term infection state such as pseudolysogeny ^9^.

Clustering analyses of 22 phages with a minimum genome size of 100 kbp (including the two outgroup genomes) based on presence/absence of protein families indicated they shared a total of 16 protein families (**Supplementary Figure 8, Supplementary Table 7**). Phosphate starvation-inducible protein PhoH (“fam582”) was the only predicted protein detected in all 22 phages (excluding the shared predicted proteins in the conserved rpS21-encoding region described above). Other common protein families include those related to DNA replication (e.g., DNA primase/helicase, DNA polymerase, HNH endonuclease, thymidylate synthase (EC:2.1.1.45), deoxyuridine 5’-triphosphate nucleotidohydrolase (EC:3.6.1.23)), those associated with virion assembly (e.g., a phage tail sheath protein, phage baseplate assembly protein W), and those for other functions (e.g., chaperone ATPase, alpha-amylase, DegT/DnrJ/EryC1/StrS aminotransferase).

### Temporal and spatial distribution and activity of bS21-encoding phages in Lake Rotsee

To reveal the spatial and temporal distribution of the bS21-encoding phages, we focused on the Lake Rotsee data and profiled phage occurence based on the sequencing depth in the metagenomic datasets. In the water column, the bS21 encoding phages were readily detected in oxic samples, especially in the under ice samples when the whole water column was oxic (**Figure 5a**).

**Figure 5.**
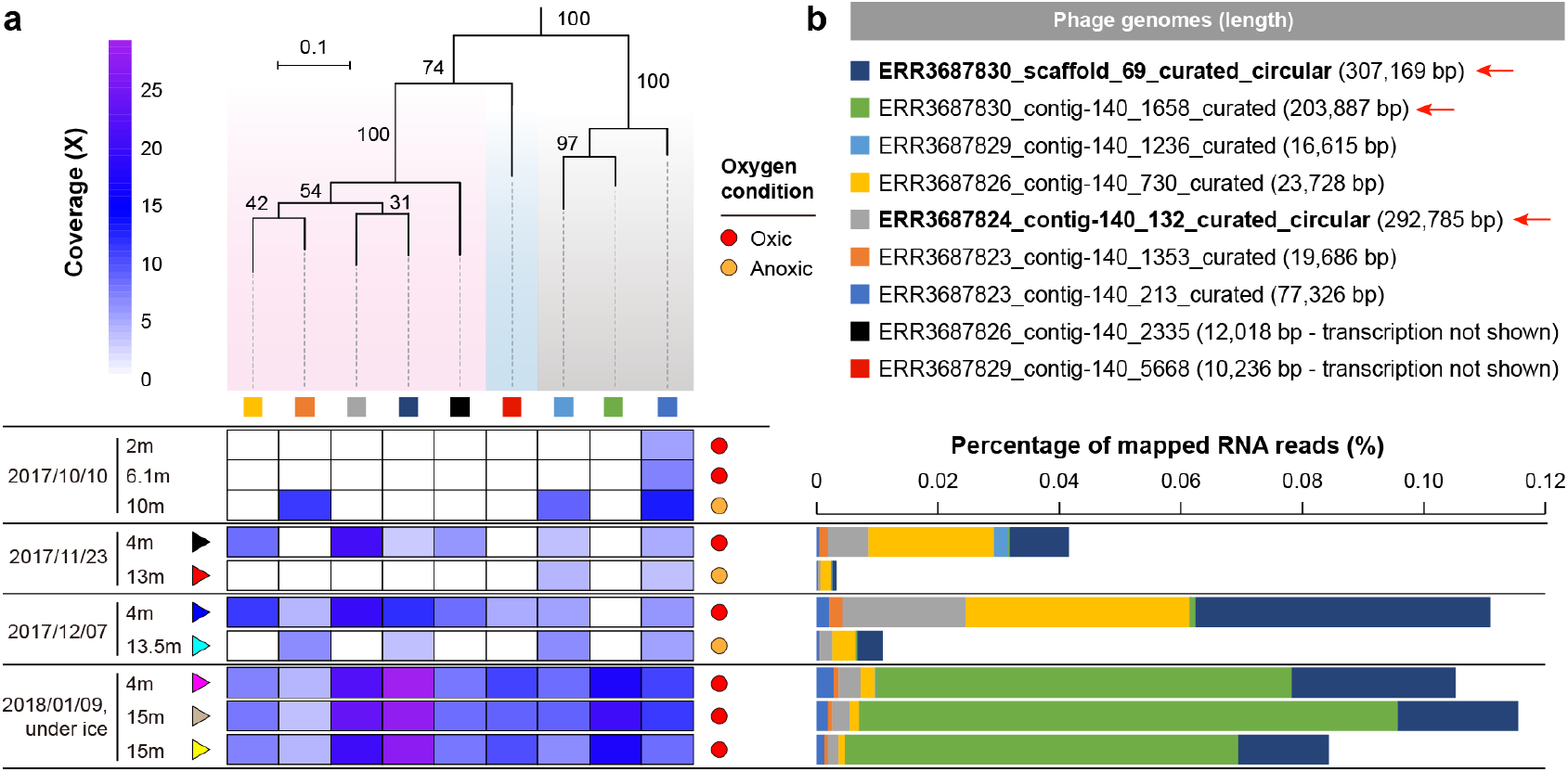
The spatial and temporal distribution and activity of bS21 phages at Lake Rotsee. (a) The sequencing coverage of each phage genome in each metagenomic dataset is shown in the heatmaps. The phages are phylogenetically clustered based on their TerL protein sequences (bootstraps shown in numbers), the colored backgrounds are the same as shown in **Figure 2** for different clades. The sampling time points and depths are shown on the left, and the oxygen conditions are indicated by colored circles on the right. Two replicates were sequenced from the 15 m sample collected in 2018. (b) The percentage of mapped RNA reads to the phage genomes in the corresponding samples (rows labeled in (a)). The mapped RNA reads had a minimum similarity of 98% to the phage genomes. No RNA data was generated for the three samples collected on 2017/10/10. The circular genomes have names in bold font.

Rotsee Lake RNA reads were mapped to the phage genomes curated from this site to reveal the transcriptional activities of bS21-encoding phages (**Figure 5b**). In general, the phages were likely to be most transcriptionally active in the oxic water columns. A total of 736 genes were transcribed in at least one sample (**Supplementary Table 8**), those for MCP, an AAA ATPase, tail sheath protein, bS21, FKBP-type peptidyl-prolyl cis-trans isomerase and a methyltransferase FkbM domain protein are among the top 100 most highly transcribed. The high transcriptional activities of MCP in five phages indicated they were in the late-stage of replication at the time of sampling.

### The transcriptional behavior of phage bS21 genes

To seek evidence of a transcriptional relationship involving bS21 and other genes we focused on the three phages that were most active based on the transcriptional level of their 19 shared single copy genes (**Figure 6a**). bS21 had very similar (but slightly lower) transcriptional activities as a neighbouring gene (hereafter, bS21_CN gene) encoded on the opposite strand. The bS21_CN gene encodes a hypothetical protein (protein family: fam498) and was not detected in the two outgroup phages without bS21 (**Supplementary Table 6**). Interestingly, comparison of the phylogenies of bS21 and bS21_CN showed a very similar evolutionary pattern (**Supplementary Figure 9**), suggesting their close functional relationship in the bS21-encoding phages.

**Figure 6.**
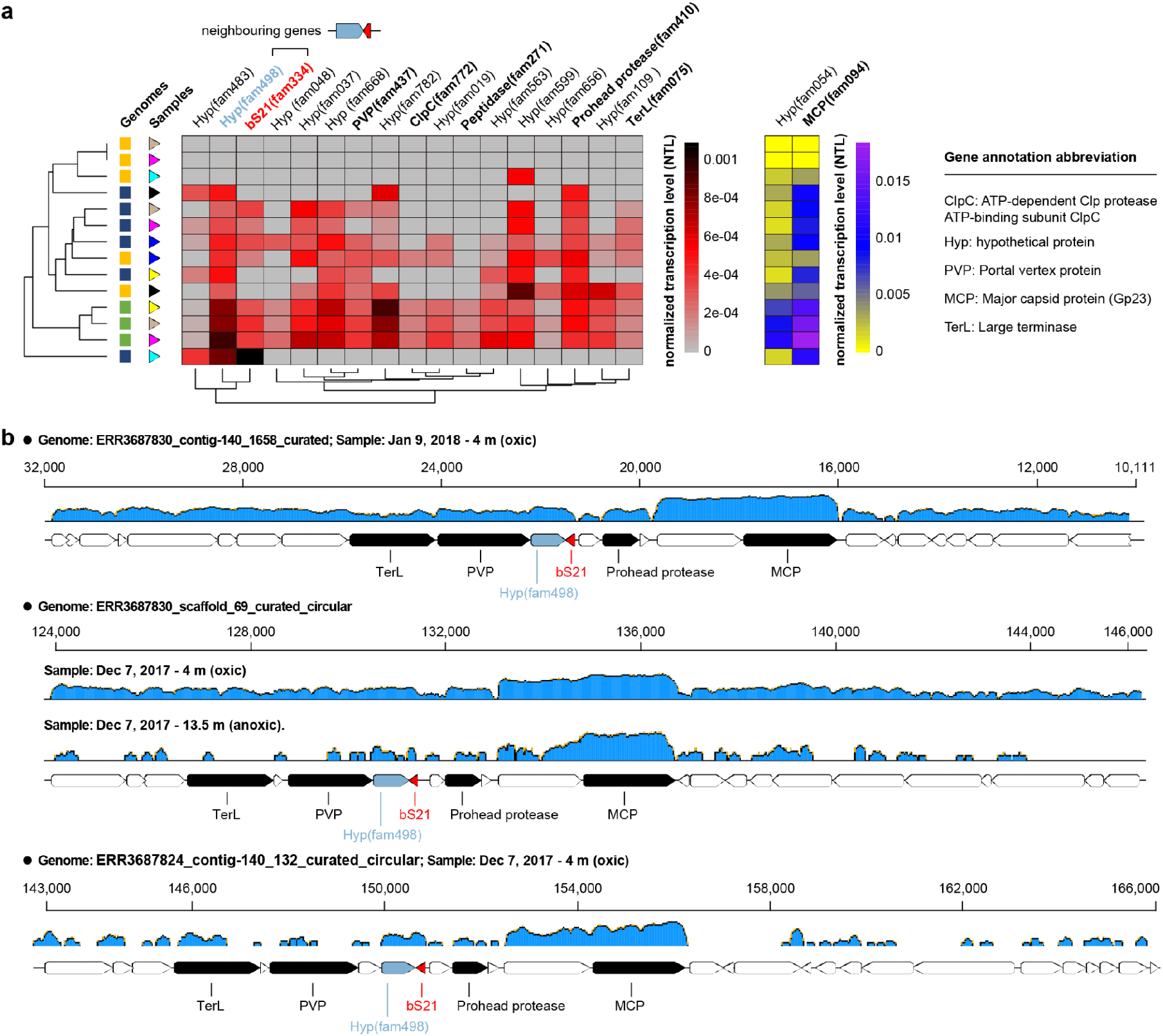
The transcription levels of bS21 and core structural protein genes. (a) The normalized transcriptional level (NTL) of shared single-copy protein families of three phages (indicated by arrows in **Figure 5b**) with ≥ 1000 RNA reads mapped. Two families (including MCP) are listed on a different scale due to their much higher transcription levels. Refer to **Figure 5** for shape symbols that designate phage genomes and samples. (b) Examples of RNA mapping profiles indicating the co-transcription of some genes neighboring bS21. Hypothetical protein genes are shown in white.

Inspection of the RNA reads mapping profiles indicated that the conserved region encoding bS21 and core structural proteins was not transcribed as an operon, whereas bS21 and bS21_CN, MCP and its upstream hypothetical protein gene, and prohead protease and its downstream hypothetical protein gene may each be transcribed together (**Figure 6b**). Given the observed RNA expression patterns, we conclude that the phage-encoded bS21 genes were actively transcribed during late-stage replication, along with other core structural proteins.

### Genomic context of bS21 genes in published phage genomes

To determine whether the phage bS21 genes are generally co-located with those for core structural proteins in diverse phages, we profiled the genomic context of bS21 in 900 published bS21-encoding phages ^2,10^ (**Supplementary Table 9**). Functional annotations were performed for the up- and down-stream 10 genes of the bS21 genes using pVOG (**Supplementary Table 10**). Of the 20 most abundant pVOGs, six were related to core structural assembly (**Figure 7a**), i.e., prohead protease (n = 310), major capsid protein (n = 154), portal vertex protein (n = 120), large terminase (n = 78), neck protein (n = 70), and a tail sheath protein (n = 29). A total of 388 genomes contained at least one of these genes within 10 genes of bS21, and eight had all of these 6 core structural proteins in close proximity. Three pVOGs were related to DNA processing, i.e., an exonuclease (n =37), an endonuclease (n = 32), DNA helicase (n = 30). Other pVOGs included Hsp20 heat shock protein (n = 127), two ATP-dependent CLP proteases (n = 50 and 47, respectively), and lysozyme (for lysis; n = 29). Interestingly, the prohead protease and the major capsid protein pVOG genes are very close to the bS21 gene (generally 2-4 genes; **Figure 7b**), as in the bS21-encoding phage genomes analyzed in this study (2-6 genes away; **Figure 1, Supplementary Figure 1**).

**Figure 7.**
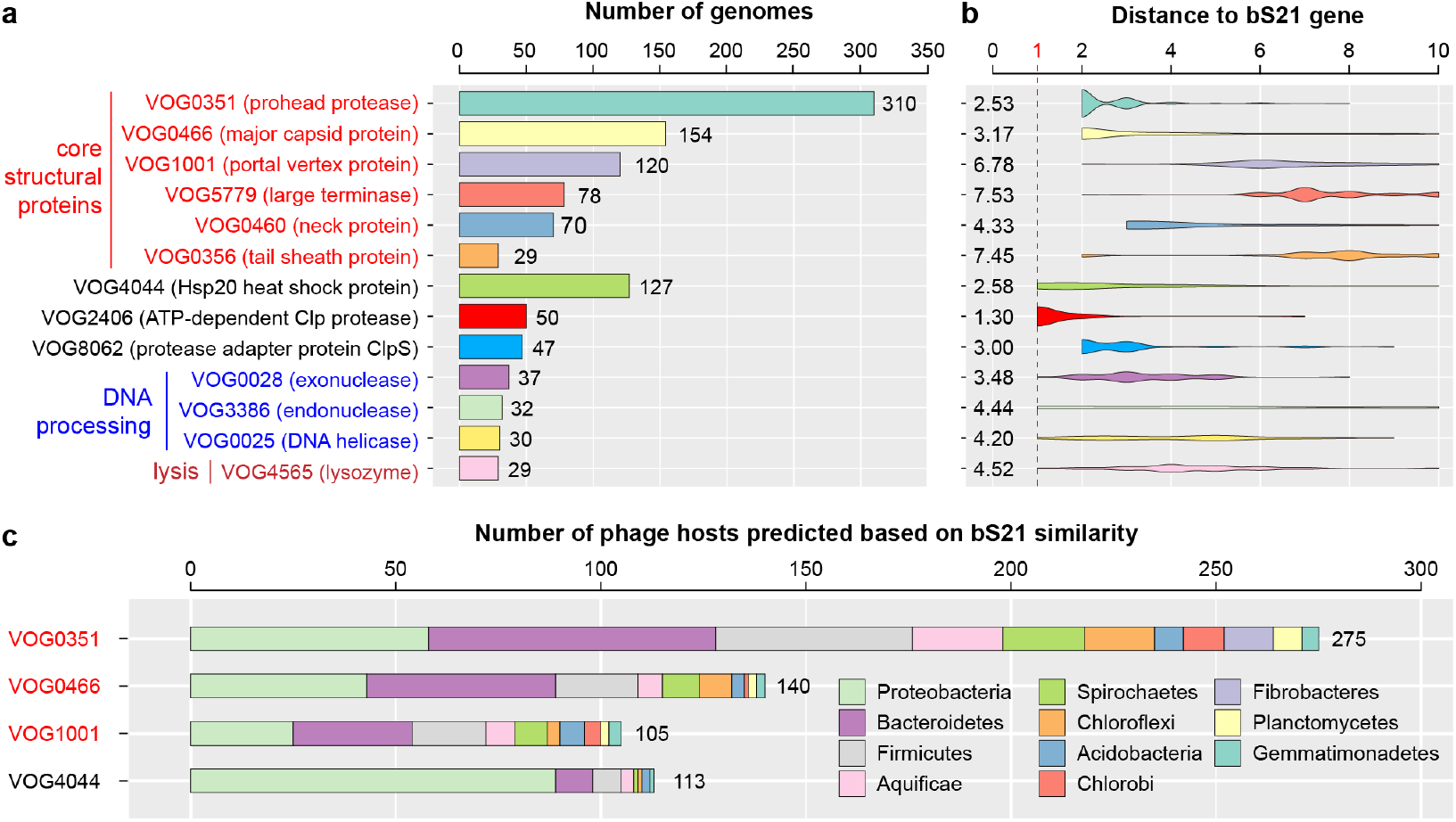
Neighbouring genes within 10 genes of bS21 in published bS21-encoding phage genomes. (a) The annotation and corresponding functional category (if assigned) of the 20 most commonly detected pVOG genes and their predicted functions are shown on the left, the total number of genomes with the gene are shown on the right. (b) The distribution of distance of each gene to bS21 in the genomes. The position of genes next to bS21 (thus distance = 1) is highlighted using a red dashed line. The average distance of each gene to bS21 is shown on the left. (c) The predicted hosts of bS21-encoding phages with the top 4 most abundant genes detected within 10 genes of bS21. The total count of hosts is shown on the right.

We respectively predicted the hosts of the bS21-encoding phages with the four most dominant pVOGs within 10 genes of bS21 (**Figure 7c, Supplementary Table 11**),. The bacterial hosts are diverse, and include Proteobacteria, Bacteroidetes and Firmicutes.

## Discussion

### The Bacteroidetes-infecting bS21-encoding phages are abundant and active in oxic water columns

Bacteroidetes, including Bacteroidia or Flavobacteriia, were ubiquitous in the analyzed samples and are the predicted hosts of most of the newly reported bS21-encoding phages (**Figure 3**). Bacteroidia spp. are strictly anaerobic ^11^ whereas Flavobacteriia spp. are strictly aerobic ^12,13^, in line with the general detection of Bacteroidia in anoxic samples and Flavobacteriia spp. in oxic samples (**Figure 3**). The majority (51/61) of the phage-encoding bS21 genes in this study are most similar to those from Flavobacteriia (**Supplementary Table 5**), explaining why most of the bS21 phages, and the most transcriptionally active subset, were detected in the oxic water column. Flavobacteriia spp. likely degrade high molecular weight compounds such as polysaccharides and proteins ^12,14^. Based on the detection of phage genes for these functions (**Supplementary Table 6**), we conclude that the bS21-encoding phages may impact carbon and nitrogen cycling in oxic water columns, both by predation of their hosts and through augmenting their metabolic capacities.

### Features indicating takeover of bacterial ribosomes by bS21-encoding phages

Some highly similar phages detected in lakes separated by thousands miles (for example, **Supplementary Figure 2b**), share identical bS21 genes in conserved genomic context, despite sequence divergence throughout the rest of the genome. This points to the high functional importance of bS21 in the phages. The conservation of the C termini of phage bS21 proteins across all of the phage clades that we defined using TerL phylogeny (**Supplementary Figure 8**) may indicate that the phage bS21 C termini are important for the phage proteins to substitute for the bacterial bS21 in the ribosomes.

bS21 is composed of two α-helices that interact in different ways with the 16S rRNA. The N-terminal α-helix is situated on top of the ASD where the mRNA would bind, whereas the C-terminal α-helix is tucked between the ASD and the rest of the 16S rRNA (**Figures 4a**) and anchors bS21 to the 16S rRNA ^3^. Our results are congruent with these observations, since sequences of phage bS21 show strong divergences from the bacterial sequences in the N-terminal α-helix (**Figures 4b**). These changes should not alter the binding of bS21 to the 16S rRNA but may provide specificity for attracting phage-specific mRNA.

All bS21 phages had either Tyr54 (54 out of 61) as occurs in *Flavobacteriia johnsoniae,* or a residue with an aromatic ring (7 out of 61) near the C-terminus of bS21 (**Supplementary Figure 6**). It has been suggested that this helps to block the ASD sequences ^8^. Thus, the phage bS21 may function in essentially the same way as that of their Bacteroidetes hosts. We infer that once phage bS21 proteins are available, the bS21 incorporates into the bacterial ribosomes, potentially enabling the phages to have their mRNA transcripts translated preferentially over the host transcripts.

### Why might phages use bS21 to hijack the ribosome in the late-stage of replication?

Our RNA analyses showed the simultaneous transcription of the genes for bS21 and core structural proteins (e.g., capsid proteins, prohead protease, large terminase, scaffolding proteins, and tail proteins; **Figure 6, Supplementary Table 8**), suggesting the potential significance of bS21 in the late-stage replication ^15^. The genomic placement and timing of transcription of bS21 genes makes sense given the huge number of proteins needed for assembly and packaging. On the other hand, a previous study showed that stalling of phage protein synthesis is an important defense strategy for a Bacteroidetes species, i.e., *Cellulophaga baltica* (class Flavobacteriia; ^16^). The replacement of bacterial bS21 with phage bS21 may be a mechanism that counters this defense strategy.

Our genetic context analyses of published bS21-encoding phage genomes showed that many bS21 genes are co-located with genes for core structural proteins, and sometimes with genes for DNA replication and lysis. Thus, we suggest that the acquisition and timing of expression of bS21 may be a more general and consistently evolved phenomenon across diverse phage lineages. This motivates future analyses that could experimentally investigate phage-encoded bS21, especially when the genes co-occur with DNA processing or lysis genes, and test the hypothesis that bS21 may be important for efficient translation of the nearby genes.

## Conclusion

By carefully manual curating 9 huge phage genomes to completion, we accurately determined genome sizes, genome organization and gene inventories (e.g., lack of other genes encoding ribosomal proteins). Partial curation further constrained the genome sizes of other phages and ensured that all key protein sequences are correct. Given RNA expression of bS21 and the flanking structural proteins in several transcriptionally active phages, we suggest that the bS21 genes in phages that infect some freshwater Bacteroidetes species are important in the late-stage replication. Our analyses of publicly available genomes suggest that this phenomenon may be more general across phage groups that infect diverse bacterial hosts.

## Materials and methods

### A global search for related phage sequences encoding bS21

To detect related phage sequences encoding the detected bS21 gene, a BLASTp search using the large terminase (TerL) protein was performed against all metagenomic datasets in ggKbase (ggkbase.berkeley.edu). All the contigs/scaffolds bearing the top BLASTp top hits were manually checked for bS21. It was confirmed that when the TerL similarity dropped to below ^~^70% similarity, the bS21 that located near the TerL was no longer observed. Two of the scaffolds with the highest TerL similarity to that of bS21 containing contigs/scaffolds but without bS21 were included in this study as an outgroup for phylogenetic and comparative genomic analyses.

### Manual curation of phage genomes

To ensure the phage genomes reported in this study were without any assembly errors, manual curation was conducted on each of them individually as described previously ^17^. Firstly, the corresponding metagenomic quality paired-end reads were mapped to the scaffold using bowtie2 ^18^ with default parameters, and the sam file was filtered to remove unmapped pairs using shrinksam (https://github.com/bcthomas/shrinksam). The shrunk sam file was imported into Geneious Prime version 2021.1.1 (https://www.geneious.com). The manual check was performed to identify any assembly errors (regions lacking paired read support), which were subsequently fixed using unplaced paired reads as previously described ^17^. Those scaffolds with sufficient coverage (generally ≥ 20X) were selected for curation to completeness. For all of the curated scaffolds, the paired high quality reads were re-mapped to the genome sequences for a final check. Regions that could not be covered by reads are scaffolding gaps and indicated by 10 Ns.

### Retrievement of related phage genomes from IMG/VR

To reveal the distribution of related phages encoding bS21 in public databases, the IMG/VR database (version 2020-10-12_5.1) was downloaded ^19^. The IMG/VR proteins were searched against the TerL protein sequences predicted from the curated genomes (see above) using BLASTp. The BLASTp hits with a minimum similarity of 97% and longer than 400 aa in length were filtered for manual inspection to include only those from IMG datasets that have been published. As a result, a total of 14 IMG/VR genomes from three sampling sites ^20,21^ which are highly similar to our bS21-encoding phages were included in our analyses (**Supplementary Table 1**), which were determined to be public by checking “published” on IMG/VR website for usage.

### Phylogenetic analyses

To reveal the phylogenetic relatedness of the phages, phylogenetic trees were built using TerL. The protein sequences were aligned using MUSCLE ^22^ with default parameters and filtered using trimal-trimAl v1.4.rev15 ^23^ to remove columns comprising ≥ 90% gaps. The trees were built by IQtree version 1.6.12 ^24^ with 1000 bootstraps using the “LG+G4” model. For the phylogeny of bS21, the alignment, filtering and tree construction of bacterial and phage bS21 protein sequences were performed the same as did for the TerL sequences.

### Comparative analysis between phage and bacterial bS21

To understand the differences between phage and bacteria encoded bS21 we built a sequence alignment of phage bS21 sequences, most closed bacterial bS21 sequences in the corresponding metagenomic samples, and publicly available bacterial bS21 sequences. The alignment was built using MUSCLE v3.8.31 with default parameters ^22^. Zebra2 ^25^ was used to perform a search for divergent positions in the generated alignment. Results were mapped on the bS21 structure from the Bacteroidetes representative *F. johnsoniae* (PDB ID: 7JIL) ^8^.

### Genomic context analysis

Protein sequences for the combined phage set were predicted using Prodigal version 2.6.3 using the “-m” model ^26^. To investigate the genomic context of bS21 in phage genomes, we gathered protein sequences within a 10 open reading frame (ORF) distance (or, to the scaffold end) in both genomic directions of the identified bS21 genes. Each ORF was assigned a genomic position relative to the bS21 (position 0). All ‘neighboring’ proteins were subjected to a two-part, de novo protein clustering pipeline in which proteins are first clustered into “subfamilies” and highly similar/overlapping subfamilies are merged using and HMM-HMM comparison approach (--coverage 0.75) (Méheust et al. 2019). We next searched all neighboring proteins against Pfam (https://pfam.xfam.org) and pVOG (https://dmk-brain.ecn.uiowa.edu/pVOGs) ^27^ HMM collections and retained hits with e-value < 1e-5. Consensus annotations for each family were obtained by computing the HMM with the most above-threshold hits among member sequences of the family (minimum 5% of member sequences). If no HMMs met these thresholds, the protein family was labeled “hypothetical” (hyp). Finally, we plotted the frequency at which each protein family was found as a function of its relative position to the focal bS21 gene (**Figure 1**). Additionally, we plotted genomic diagrams of individual regions of interest using gggenes (https://wilkox.org/gggenes).

### Microbial community composition analyses

To reveal the community composition of the samples analyzed in this study, the protein-coding genes were predicted using Prodigal ^26^ (-m = meta) from all metagenomic assembled sequences with a minimum length of 1000 bp. The ribosomal protein S3 (rpS3) was predicted using hmmsearch (version HMMER 3.3) ^28^ with the hmm database from TIGRFAM ^29^. For taxonomic information, the predicted rpS3 protein sequences (minimum length, 100 aa) were searched using BLASTp against the rpS3 proteins from NCBI RefSeq and those of Candidate Phyla Radiation reported previously ^30^, and taxonomy of the best hits was used. The nucleotide sequences of the predicted rpS3 genes (minimum length, 300 nt) were clustered using cd-hit-est (parameters: -c = 0.97, -aS = 0.5, -aL = 0.5, -G = 1) ^31^ to generate a non-redundant dataset. The quality metagenomic reads from each sample were individually mapped to the non-redundant dataset and filtered allowing ≤ 3% mismatch. The coverage of each rpS3 sequence was determined by the total mapped bases dividing by its length.

### Host prediction of phages

To predict the bacterial hosts of the phages, CRISPR-Cas spacers were searched for targeting. Firstly, all the scaffolds with a minimum length of 1 kbp from all the corresponding samples with the bS21 phages were predicted for CRISPR repeat arrays using PILER-CR ^32^ with default parameters. The protein-coding genes within 10 kbp of both up- and down-stream of each repeat array were predicted and searched for Cas proteins using the TIGRFAM HMM database ^29^ with hmmsearch (version HMMER 3.3) ^28^. For the scaffolds with both CRISPR repeat arrays and at least one cas protein, spacer sequences were extracted. Spacers were also extracted from the mapped reads and unplaced paired reads that may carry divergent spacers. The extracted spacers were searched against the manually curated phage sequences using blastn-short ^33^ with parameters as follows: -evalue = 1e-3, -perc_identity = 70. The search results were parsed to retain those hits with a minimum match of 24 nt and no more than one mismatch ^2^, or 30 bp with no more than 3 mismatches. The phages whose scaffolds had matches to the spacers were considered as the likely bacterial hosts.

### The temporal and spatial distribution of bS21 phages by read mapping

The samples with bS21 phage genomes reconstructed in this study and their related samples were mapped to all the curated bS21 phage genomes using Bowtie2 ^18^. A given bS21 phage was considered to be detected in a given sample if 90% of its genome was covered with at least one read (minimum nucleotide sequence similarity of 97%), and the coverage of this phage in the sample was accordingly determined as the total length of mapped reads dividing by the total covered genome length. A custom python script (https://dCov.py) was prepared to perform the analyses.

### Gene annotation and metabolic prediction

The tRNAs encoded on all the curated phage genomes and retrieved IMG/VR sequences were predicted using tRNAscanSE (version 2.0.3) ^34^. The predicted protein-coding genes were annotated by searching against the databases of Kyoto Encyclopedia of Genes and Genomes (KEGG) ^35^, UniRef100 ^36^ and UniProt ^37^ using Usearch (version v10.0.240_i86linux64) ^38^ with an e-value threshold of 10^−4^. For specific metabolic potential of interest, the predicted protein-coding genes were also investigated using online HMM search tools.

### Metatranscriptomic analyses

The seven raw metatranscriptomic RNA datasets from Lake Rotsee were downloaded ^39^ and filtered to remove sequencing contamination, adaptors and low quality bases/reads as performed for metagenomic reads (see above). To profile the transcription of protein-coding genes, the quality RNA reads were mapped to the curated phage genomes reconstructed from Lake Rotsee using Bowtie2 ^18^ allowing 3 mismatches each read (i.e., 98% similarity). The normalized transcriptional level of a given protein-coding gene (*gene_a*) in a given sample was determined by calculating as follows: total_base_*gene_a*_/length_*gene_a*_/total_read_*genome_a*_, in which total_base_*gene_a*_ means the total bases mapped to a given gene, length_*gene_a*_ means the length of the nucleotide sequence of the gene, and total_read_*genome_a*_ means the total number of reads mapped to the corresponding genome. Only those protein-coding genes with at least 80% of the bases covered were calculated for normalized transcriptional level.

### Genomic context of bS21 in published bS21-encoding phage genomes

To reveal if the bS21 genes in the published viral genomes are also co-located with these for core structural proteins, we checked all the huge phages genomes reported by Al-Shayeb et al. ^2^, and also the viral genomes reconstructed from the Global Ocean Virome (GOV) ^10^. The protein-coding genes were searched against the bS21 HMM from TIGRFAM ^29^ using hmmsearch (version HMMER 3.3) ^28^ with the parameters of “--cut_tc”. The results were parsed using cath-resolve-hits ^40^. We identified bS21 genes in 68 huge phages ^2^ and 832 GOV viral genomes. The genomic context of the bS21 genes in these 900 genomes were performed as described above (see section “**Genomic context analysis**”). For the phage bS21 genes with genes for some specific proteins within 10 genes, the taxonomy of their most similar bacterial bS21 were evaluated by comparison against the NCBI RefSeq bS21 proteins using BLASTp ^33^, the hits with the highest bit scores were retained for further analyses. When we checked the IMG/VR genomes for bS21 there were more than 14,000 bS21 hits. However, given the policy of IMG data use, we restricted our analyses to the published genomes.

## Accession numbers

The genomes of the bS21-encoding and outgroup phages have been deposited at NCBI under BioProject xxx (TBA), and also available from ggkbase https://ggkbase.berkeley.edu/PS21/organisms (please sign in by providing your email address to download) and at figshare (https://figshare.com/articles/dataset/bS21_encoding_phages/16744504).

## Acknowledgements

The study was supported by the Moore Foundation Grant 71785. We also thank the Chan Zuckerberg Biohub and the Innovative Genomics Institute at University of California, Berkeley for funding support. We thank Anne-Catherine Lehours, Cindy Castelle, Corinne Bardot, Hermine Billard, Jonathan Colombet, and Fanny Perriere for assistance collecting and preparing the Lac Pavin samples. We acknowledge the researchers that generated the public data used in this study, especially Magdalena J. Mayr et al. who published the Lake Rotsee data.

## Author contributions

The study was initiated by J.F.B.. Manual genome curation was performed by L.X.C. and J.F.B.. Genomic context and protein family analyses were conducted by A.L.J., A.L.B. and L.X.C.. Comparative genomic analyses, microbial composition, host-phage relationship determination, phylogenetic analyses, tRNA analyses were performed by L.X.C.. The distribution of phages in samples by read mapping was conducted by L.X.C. and A.L.J.. Metabolic potential and metatranscriptomic analyses were conducted by L.X.C. and A.L.B.. The bS21 related analyses were performed by L.X.C. and P.I.P.. T.C.N. and L.A.W. collected and prepared the Canada samples for sequencing. The manuscript was written by L.X.C. and had input from all authors.

