## Supplementary Figures 1-9 for "Phage-encoded ribosomal protein S21 expression is linked to late stage phage replication"

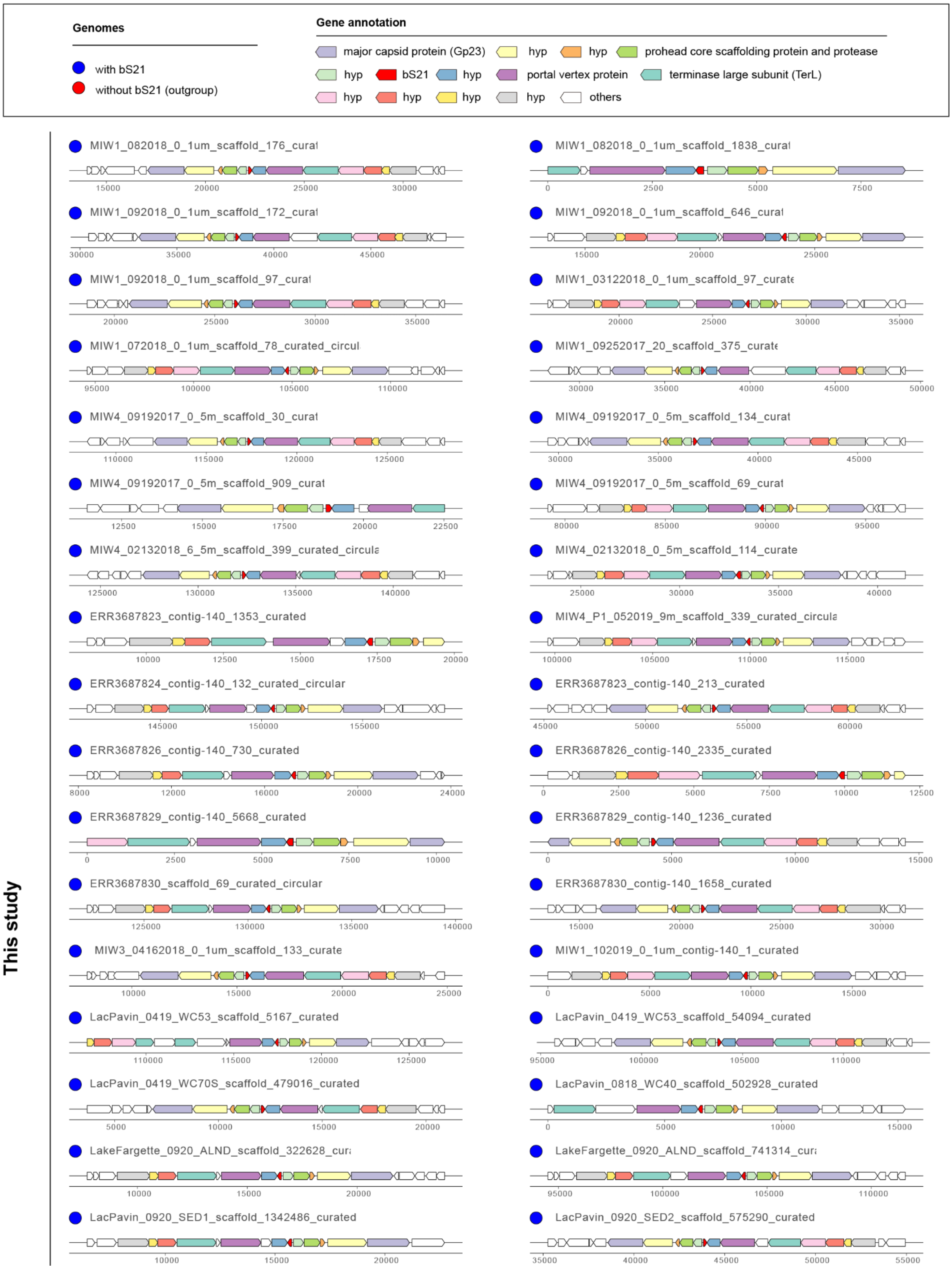
Genomes
with bS21
without bS21 (outgroup)
Gene annotation
major capsid protein (Gp23)
hyp
hyp
prohead core scaffolding protein and protease
hyp
bS21
hyp
portal vertex protein
terminase large subunit (TerL)
hyp
hyp
hyp
hyp
others
This study
MIW1_082018_0_1um_scaffold_176_curat
MIW1_082018_0_1um_scaffold_1838_curat
MIW1_092018_0_1um_scaffold_172_curat
MIW1_092018_0_1um_scaffold_646_curat
MIW1_092018_0_1um_scaffold_97_curat
MIW1_03122018_0_1um_scaffold_97_curate
MIW1_072018_0_1um_scaffold_78_curated_circul
MIW1_09252017_20_scaffold_375_curate
MIW4_09192017_0_5m_scaffold_30_curat
MIW4_09192017_0_5m_scaffold_134_curat
MIW4_09192017_0_5m_scaffold_909_curat
MIW4_09192017_0_5m_scaffold_69_curat
MIW4_02132018_6_5m_scaffold_399_curated_circula
MIW4_02132018_0_5m_scaffold_114_curate
ERR3687823_contig-140_1353_curated
MIW4_P1_052019_9m_scaffold_339_curated_circula
ERR3687824_contig-140_132_curated_circular
ERR3687823_contig-140_213_curated
ERR3687826_contig-140_730_curated
ERR3687826_contig-140_2335_curated
ERR3687829_contig-140_5668_curated
ERR3687829_contig-140_1236_curated
ERR3687830_scaffold_69_curated_circular
ERR3687830_contig-140_1658_curated
MIW3_04162018_0_1um_scaffold_133_curate
MIW1_102019_0_1um_contig-140_1_curated
LacPavin_0419_WC53_scaffold_5167_curated
LacPavin_0419_WC53_scaffold_54094_curated
LacPavin_0419_WC70S_scaffold_479016_curated
LacPavin_0818_WC40_scaffold_502928_curated
LakeFargette_0920_ALND_scaffold_322628_cur
LakeFargette_0920_ALND_scaffold_741314_cur
LacPavin_0920_SED1_scaffold_1342486_curated
LacPavin_0920_SED2_scaffold_575290_curated

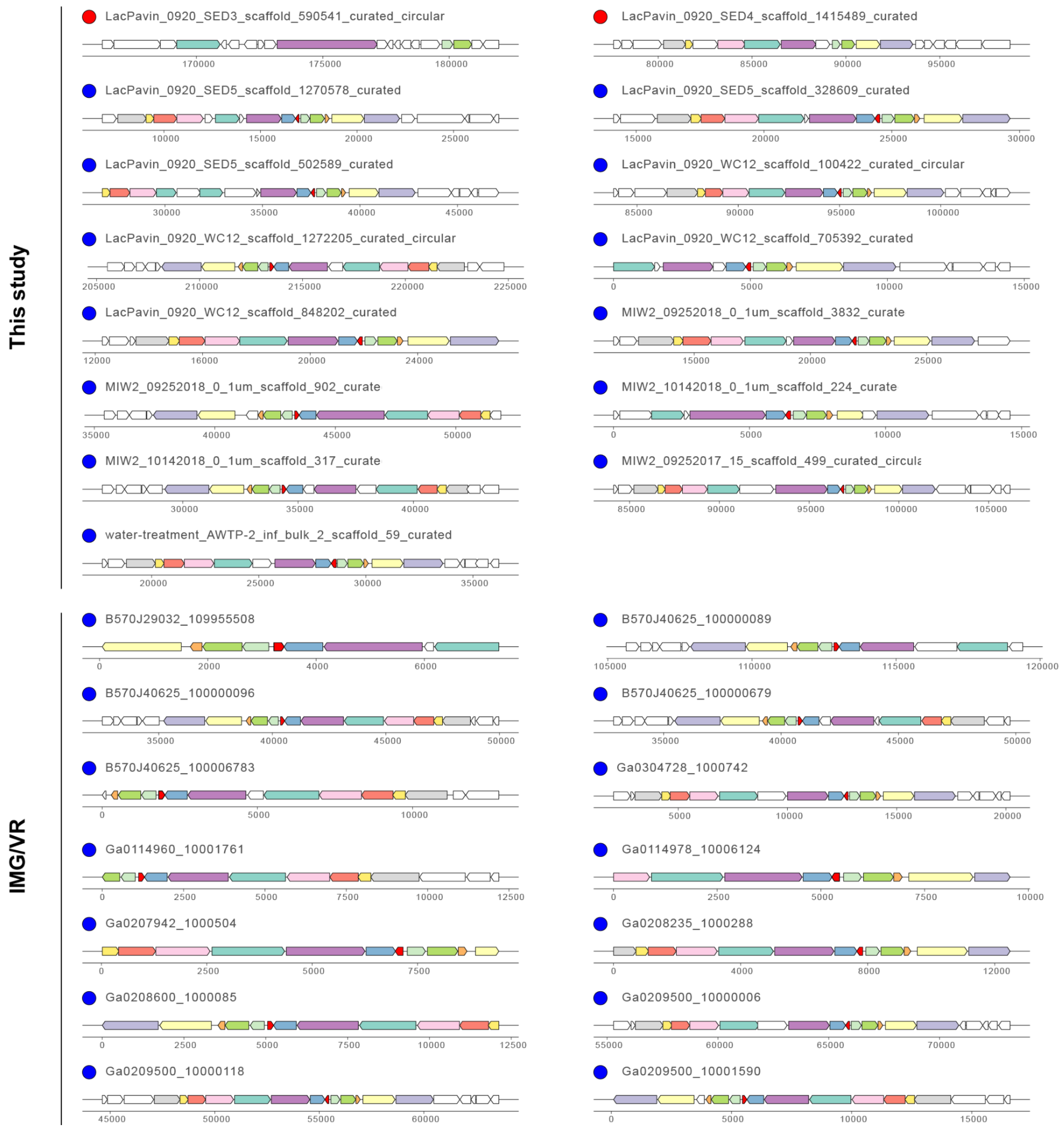


**Supplementary Figure 1 | Detailed genetic context of the bS21 genes in all the genomes reported in this study and retrieved from IMG/VR.** The 12 most frequent genes near the bS21 gene (in red) are shown in different colors (same as in **Figure 1** in the main text). For the two outgroup genomes, the similar regions are present. The scale bars and numbers under the genes indicate the corresponding positions on the scaffolds.

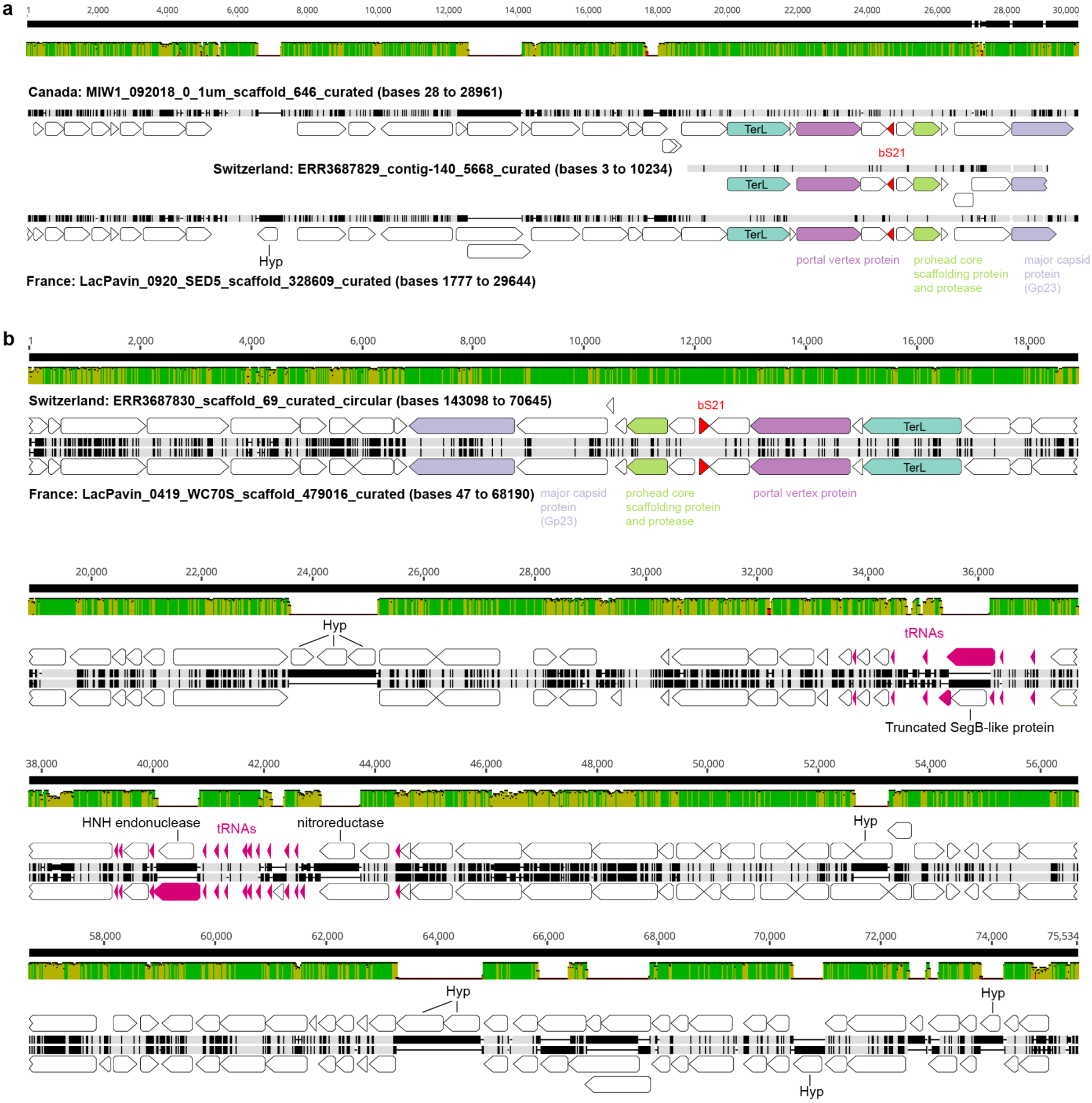


**Supplementary Figure 2 | Comparison of phage genomic fragments reconstructed from different sampling sites.** (a) Three genomes from MIW1 (Canada), Lake Rotsee (Switzerland) and Lac Pavin (France). (b) Two genomes from Lake Rotsee (Switzerland) and Lac Pavin (France). The top scale bar indicates the comparative locations, the green/gray profile shows the nucleotide sequence similarity among sequences. The aligned region for each sequence is shown in the bracket following its name. In the alignment of each sequence, black lines are for different bases and white for shared bases. The protein-coding genes with known functional annotation and shared by all are indicated by different colors, all others are shown in white. The annotation of divergent genes between genomes are shown. Hyp, hypothetical protein.

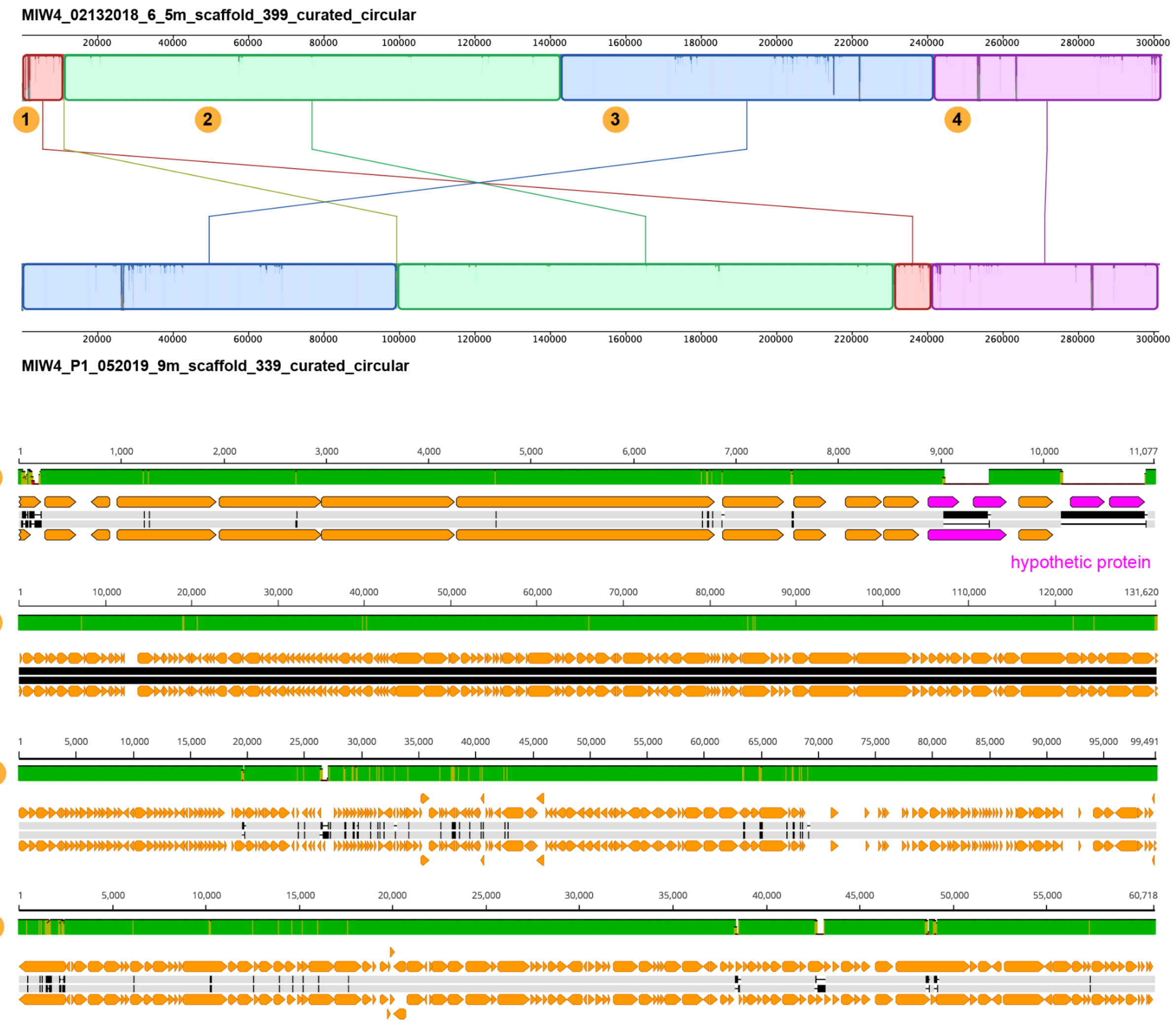


**Supplementary Figure 3 | Comparison of two complete genomes reconstructed from the same site but different time points shows variations.** Mauve genome alignment viewer is shown at the top, the details of the four regions are shown at the bottom individually. The scale bars and numbers indicate the location of each region. The green/gray profile (0-100%) under the scale bar indicates the nucleotide sequence similarity between the region of two genomes.

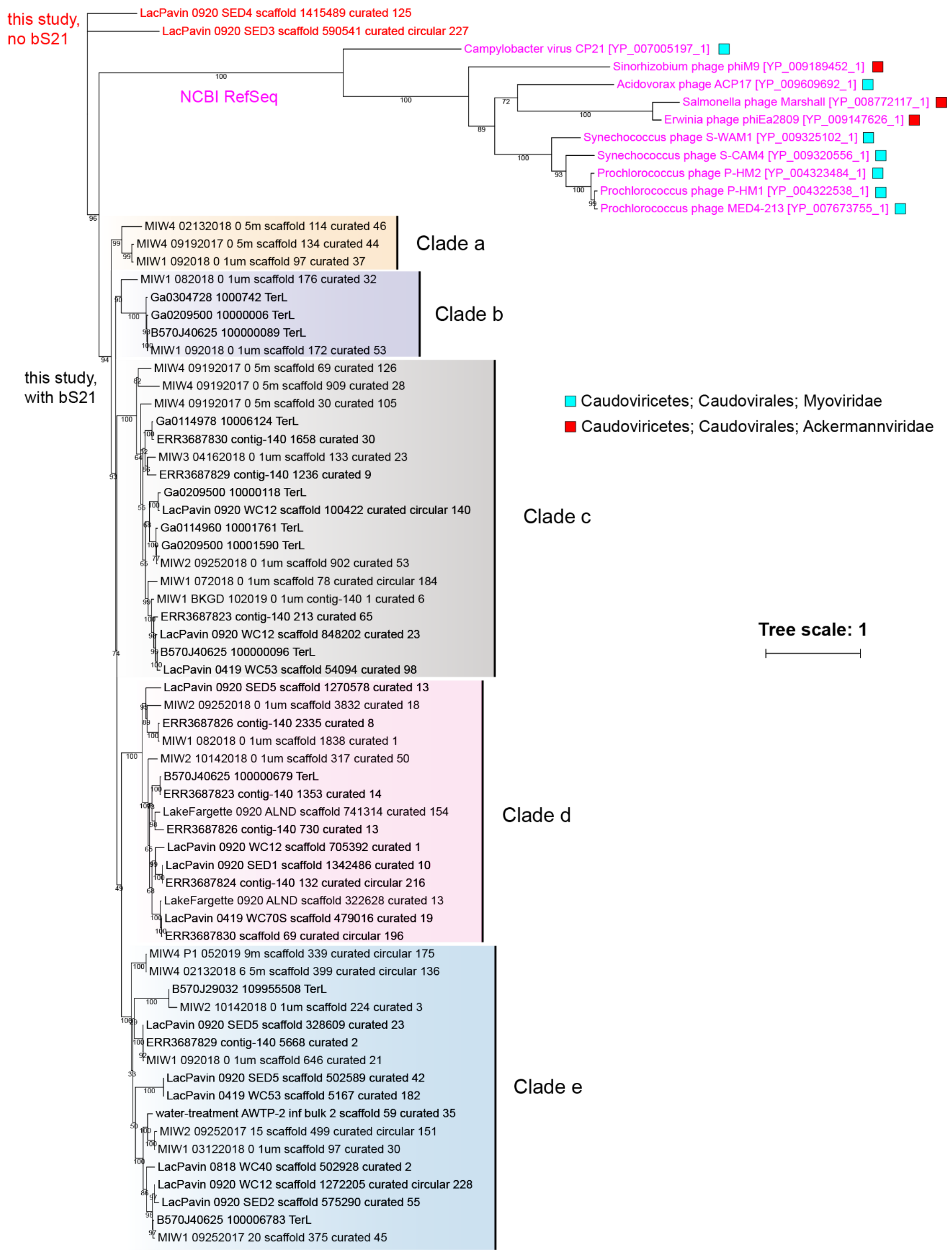


**Supplementary Figure 4 | Phylogenetic analyses of bS21-encoding phage based on TerL including references from NCBI RefSeq.** The taxonomic assignment of RefSeq viruses is indicated by colored squares. The assigned clades of bS21-encoding phages are the same as shown in **Figure 2** of the main text.

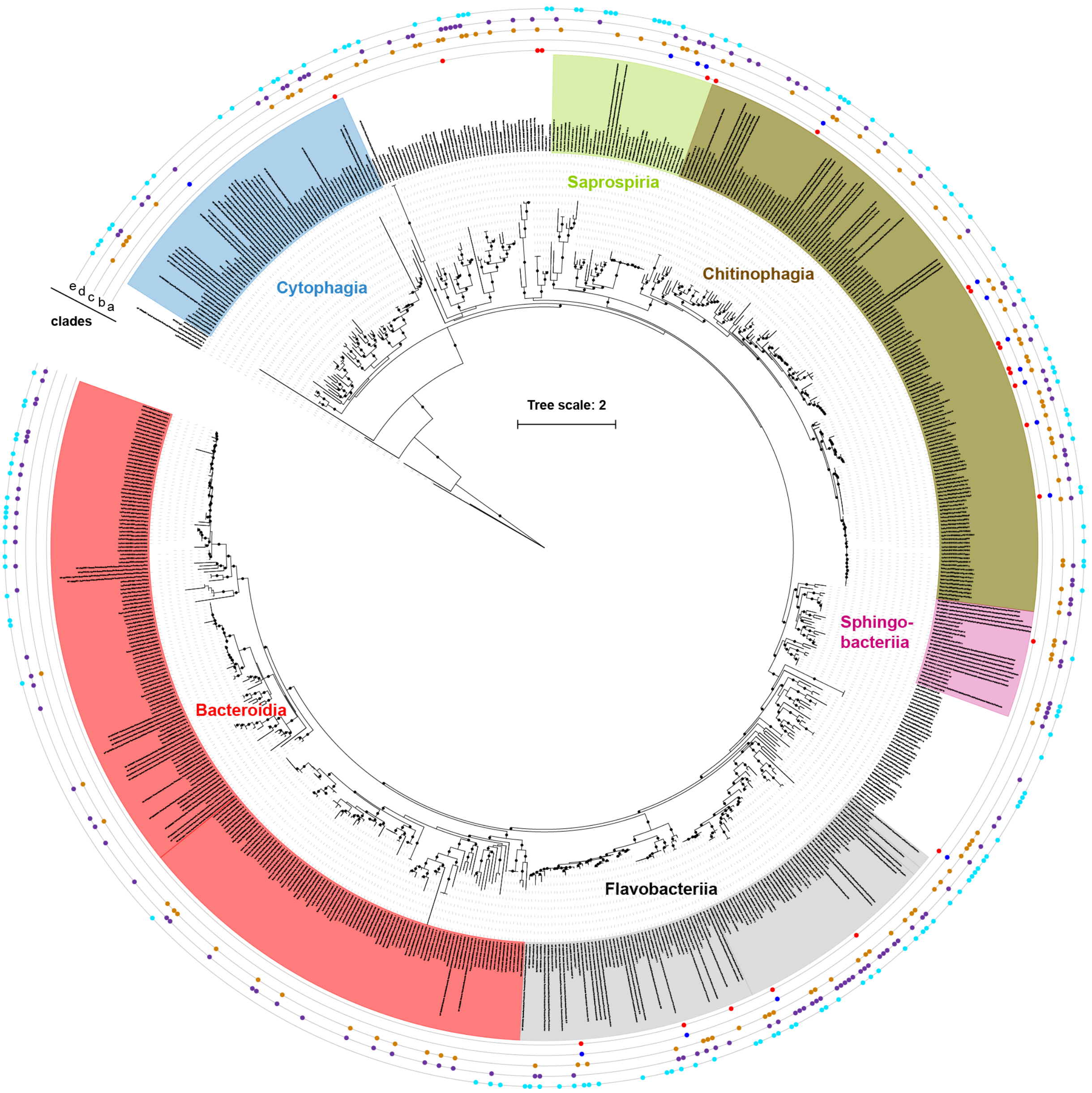


**Supplementary Figure 5 | The co-detection of bS21-encoding phage clades with Bacteroidetes members.** The tree was built including all the Bacteroidetes rpS3 identified in all 45 samples analyzed in this study and NCBI RefSeq sequences. Black circles on the tree indicate bootstrap ≥ 70. The classes of Bacteroidetes were indicated with color background if the corresponding clades were with RefSeq sequences. The detection of the five bS21-encoding phage clades was indicated by colored circles, once the phage clade was identified in the corresponding sample where the Bacteriodetes member was from.

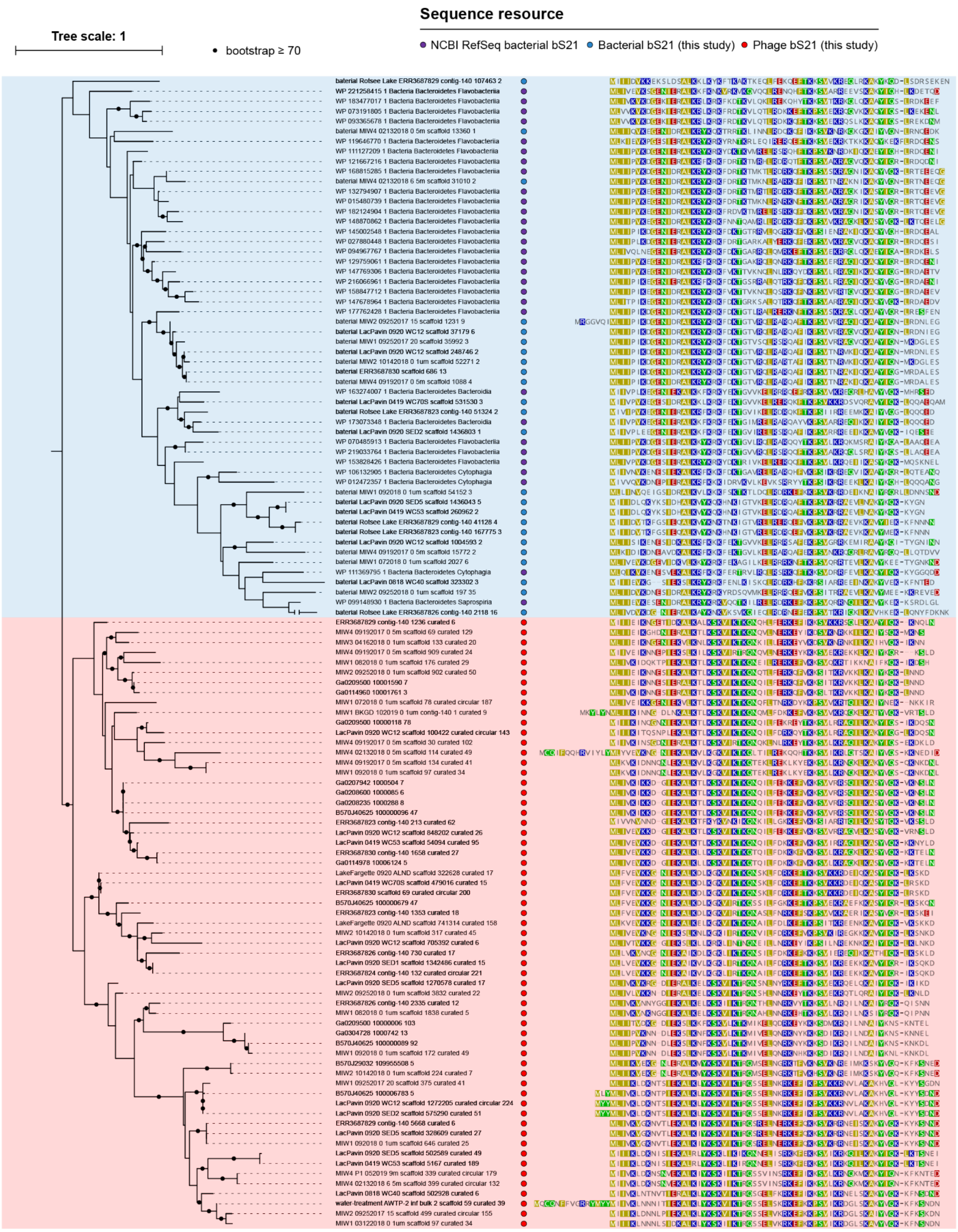


**Supplementary Figure 6 | Phylogenetic analyses and alignment of bS21 protein sequences encoded by bacteria and phages.** The resources of the sequences are indicated by colored circles. The sequence logo and consensus sequences of bacterial and phage bS21 alignments are shown at top right corner, the middle ‘core’ part alignment is also included.

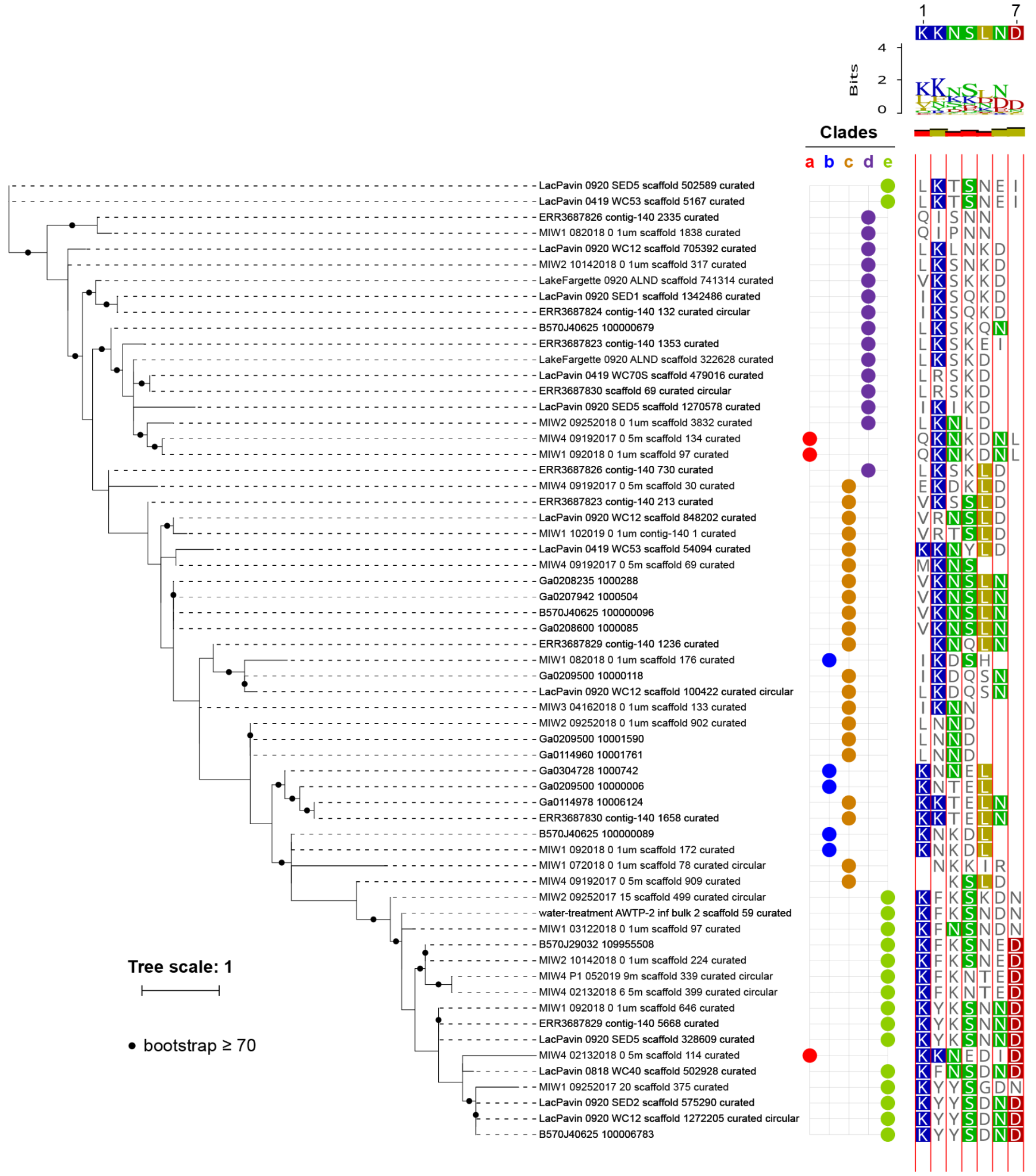


**Supplementary Figure 7 | Phylogenetic analyses and alignment of the C terminal of bS21 protein sequences encoded by bacteria and phages.** The clades of bS21-encoding phages determined based on TerL phylogeny (**Figure 2** in the main text) are indicated by colored circles. The alignment of the C terminal regions are shown on the right, with both consensus sequence and sequence logo shown at the top right corner.

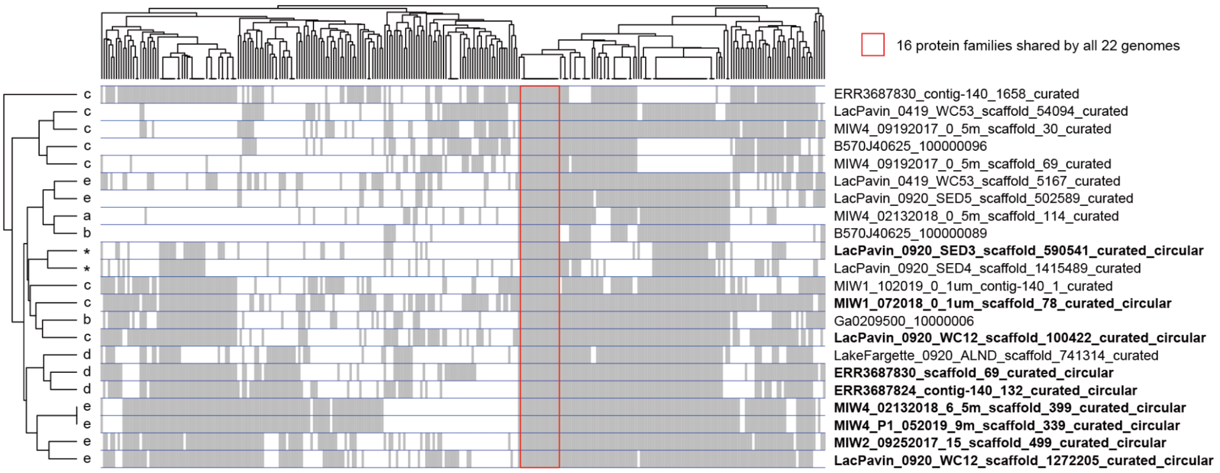


**Supplementary Figure 8 | Clustering analyses of bS21-encoding and outgroup phages.** Only those genomes with a size ≥ 100 kbp in length were included for this analysis. The clustering was based on the presence and absence of protein families detected in at least five phage genomes. The assigned clades of the phages based on TerL (**Figure 2**) are shown with letters and asterisks for the two outgroup phage genomes.

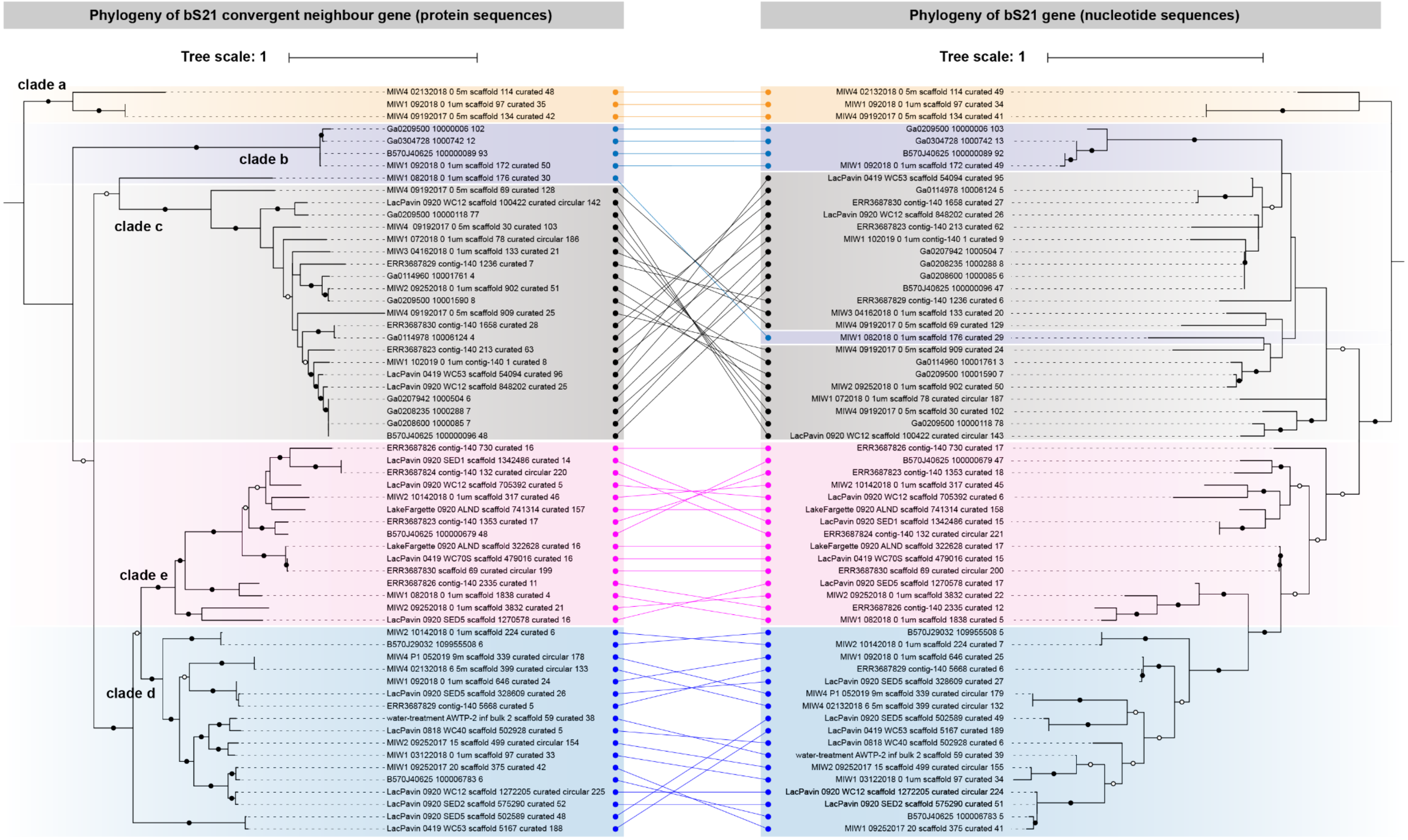


See figure legend on next page.

**Supplementary Figure 9 | Comparison of the phylogeny of bS21 (right panel) and its convergent neighbour gene (left panel).** To have a better phylogenetic resolution of the bS21 gene, the nucleotide sequences were used for analyses. The two genes from the same phage genome in the two trees were linked by lines. The phage clades determined by TerL phylogeny (**Figure 2** in the main text) are shown. A solid circle indicates bootstraps > 90, while an open circle indicates bootstraps > 79 and ≤ 90.
